# Visual field reconstruction using fMRI-based techniques

**DOI:** 10.1101/2020.07.29.226258

**Authors:** Joana Carvalho, Azzurra Invernizzi, Joana Martins, Nomdo M. Jansonius, Remco J. Renken, Frans W. Cornelissen

## Abstract

**Purpose:** To evaluate the accuracy and reliability of functional magnetic resonance imaging (fMRI)-based techniques to assess the integrity of the visual field (VF).

**Methods:** We combined fMRI and neurocomputational models, i.e conventional population receptive field (pRF) mapping and a new advanced pRF framework “micro-probing” (MP), to reconstruct the visual field representations of different cortical areas. To demonstrate their scope, both approaches were applied in healthy participants with simulated scotomas (SS) and participants with glaucoma. For the latter group we compared the VFs obtained with standard automated perimetry (SAP) and via fMRI.

**Results:** Using SS, we found that the fMRI-based techniques can detect absolute defects in VFs that are larger than 3 deg, in single participants, and based on 12 minutes of fMRI scan time. Moreover, we found that MP results in a less biased estimation of the preserved VF. In participants with glaucoma, we found that fMRI-based VF reconstruction detected VF defects with a correspondence to SAP that was decent, reflected by the positive correlation between fMRI-based sampling density and SAP-based contrast sensitivity loss (SAP) r^2^=0.44, p=0.0002.This correlation was higher for our new approach (MP) compared to that for the conventional pRF analysis.

**Conclusions:** fMRI-based reconstruction of the VF enables the evaluation of vision loss and provides useful details on the properties of the visual cortex.

**Translational Relevance:** fMRI-based VF reconstruction provides an objective alternative to detect VF defects. It may either complement SAP, or could provide VF information in patients unable to perform SAP.

## 1. Introduction

The detection of visual field defects (VFD) is an essential aspect of ophthalmic assessment. This is especially relevant in clinical pathologies such as glaucoma, for diagnosis and monitoring of disease progression^1^. Standard automated perimetry (SAP) is the clinical gold standard to measure the visual field ^2^. SAP assesses the luminance sensitivity at multiple locations of the visual field using incremental light stimuli ^3^. However, there are a number of issues that limit the applicability and reliability of SAP: SAP is fairly complicated to perform, relies on a participant’s attention and their experience, and on many other factors ^4,5,6,7^. These factors in general limit the power of SAP as a diagnostic technique, and make SAP unusable in some patients. Moreover, not all diagnostic dilemmas can be addressed easily with SAP, e.g., (suspected) functional VF loss. If we would have a technique that could accurately chart the VF without requiring participant compliance (other than lying without moving and keeping their eyes open), this would obviously benefit clinical care.

The combination of fMRI and biologically-inspired data analysis methods such as population receptive field (pRF) modeling may provide an option to realize this. Such methods model the neuronal response of populations of RFs. These approaches have become essential tools to study and assess the visual field representations in the healthy and impaired visual system ^8–10^. By back projecting the modelled population RFs, it is possible to visualize the responsiveness of a particular cortical visual area in VF coordinates. Previous studies used simulated and natural scotomas to investigate changes in visual field representations following damage to the retina ^11–18^. In particular, Hummer *et al*.^*18*^ showed that it is possible to accurately detect central scotomas that are larger than 4.7 deg (diameter). Building on this, Ritter *et al*. found decent correspondence between visual fields charted using micro-perimetry and pRF mapping in participants with central and peripheral retinal diseases ^17^. Therefore, pRF modeling holds the promise to: 1) enable visual field estimation without relying on a participant’s task performance and 2) inform about the integrity of the cortical mechanisms underlying someone’s visual field. Once available, such information could further inform on cortical reorganization, perceptual abilities, and the potential to enhance or restore vision via training or restorative therapy.

Micro-probing, a recently introduced mapping technique, enables a more accurate delineation of pRFs (Carvalho et al., 2020). Consequently, it also promises potentially more accurate charting of the VF based on fMRI. Therefore, in this study we aim to validate our new MP technique for reconstructing the VF and detecting VFDs. We do this by using simulated scotomas (SS) that mimic the lack of visual input caused by VFDs in healthy participants. In particular, we will address the capability of pRF-based techniques to detect heterogeneously shaped and sized scotomas located in different areas of the visual field. SS provides the ground truth when testing the reliability of fMRI-based approaches in detecting VFDs ^18^. In addition, SS is an essential tool in studies that investigate the plasticity of the visual cortex in patients with ophthalmic diseases. In these conditions, SS allows to control for differences in neural activity driven by differential visual input rather than neuroplastic changes ^14^. However, the presence of scotomas may result in biases in the estimated pRF properties due to partial stimulation of the underlying population of RFs ^12^. Here, we will take advantage of the different VF sampling mechanisms of MP and conventional pRF models together with SS to determine the presence of methodological biases in pRF property estimation ^11,12,19^. Ultimately, we will assess the ability to reconstruct VFD based on data obtained from different cortical areas. Finally, we will compare SAP- and fMRI-based VF maps of participants with glaucoma.

## 2. Materials and Methods

### 2.1 Participants and ethics statement

A total of 6 participants (3 females; average age: 28; age-range: 26–32 years-old) with normal or corrected to normal vision; and 19 participants with primary open angle glaucoma (10 females; average age: 70; age-range: 55–84 years-old) were recruited. Note that the normal vision participants were not meant to serve as controls to the glaucoma participants. They were used to verify the technique’s ability to detect VFDs, in a controlled manner where we knew the ground truth (the simulated scotomas). Prior to scanning, participants signed an informed consent form. Our study was approved by the University Medical Center of Groningen, Medical Ethical Committee, and conducted in accordance with the Declaration of Helsinki.

Inclusion criteria for the participants with glaucoma were as follows: having an intraocular pressure (IOP) > 21 mmHg before treatment onset, presence of a VFD (glaucoma hemifield test outside normal limits) due to glaucoma, abnormal optical coherence tomography (OCT; peripapillary retinal nerve fiber layer thickness [pRNFL] at least one clock hour with a probability of < 1 %) and spherical equivalent refraction within ±3 D.

Exclusion criteria for both groups were: having any ophthalmic disorder affecting visual acuity or VF (other than primary open angle glaucoma (POAG) in the participants with glaucoma group), any neurologic or psychiatric disorders, the presence of gross abnormalities or lesions in their magnetic resonance imaging (MRI) scans, or having any contraindication for MRI (e.g., having a pacemaker or being claustrophobic).

### 2.2 Experimental procedure

#### 2.2.1 Participants with normal vision

Each participant completed two (f)MRI sessions of approximately one hour each. In the first session, the participants were subjected to an anatomical scan and to the retinotopic mapping using luminance contrast stimulus (LCR), see Section 2.2.4 ^8^. In the second session, the participants were subjected to the retinotopic mapping with various simulated scotomas (LCR SS) superimposed. In both sessions, the participants viewed the stimuli binocularly.

#### 2.2.2 Participants with Glaucoma

##### 2.2.2.1 Ophthalmic Data

Prior to their participation in the MRI experiments, we assessed for all participants with glaucoma their visual acuity, IOP, VF sensitivity (measured using HFA and frequency doubling technology (FDT)) and retina nerve fiber layer (RNFL) thickness. Visual acuity was measured using a Snellen chart with optimal correction for the viewing distance. IOP was measured using a Tonoref noncontact tonometer (Nidek, Hiroishi, Japan). The VFs were first screened using FDT (Carl Zeiss Meditec) using the C20-1 screening mode. The contrast sensitivity at several locations of the VF was measured using SAP in particular HFA (Carl Zeiss Meditec, Jena, Germany) using the 24-2 or 30-2 grid and the Swedish Interactive Threshold Algorithm (SITA) Fast. Only reliable HFA tests were included in this study. A VF test result was considered unreliable if false-positive errors exceeded 10% or fixation losses exceed 20% and false-negative errors exceed 10% ^20^. The average fixation loss in SAP was 6% (± 10 %) and 8% (± 10 %) for the left and right eye, respectively. Table S1 shows the fixation loss, false-positive error and false-negative error obtained per participant. Finally, the RNFL thickness was measured by means of OCT using a Canon OCT-HS100 scanner (Canon, Tokyo, Japan). In this study we focused the analysis on the SAP outcomes.

##### 2.2.2.2 Neuroimaging

Each participant completed two (f)MRI sessions of approximately 1.5h each. In the first session, the anatomical scan (T1w), Diffusion Weighted Imaging (DWI), T2w, resting state functional scans and a task designed to localize the middle temporal visual area (MT) were acquired. Neurons in the MT area respond to movement, which is known to be impaired in glaucoma. The MT data was not presented in this report. In the second session, the retinotopic mapping and scotoma localizer experiments took place. These experiments were performed binocularly and monocularly as well. Here, we focused our analysis on the monocular retinotopic mapping, the most lesioned eye, was stimulated and the other was occluded using an MRI compatible opaque lense. The most lesioned eye was selected based on the SAP MD (mean deviation) score; the eye with the lowest MD was selected. The monocular retinotopy experiment comprised three runs.

### 2.3 Stimulus presentation

Stimuli were presented on an MR compatible display screen (BOLDscreen 24 LCD; Cambridge Research Systems, Cambridge, UK). The screen was located at the head-end of the MRI scanner. Participants viewed the screen through a tilted mirror attached to the head coil. Distance from the participant’s eyes to the display (measured through the mirror) was 120 cm. Screen size was 22×14 deg. Visual stimuli were created using MATLAB (Mathworks, Natick, MA, USA) and the Psychtoolbox ^21,22^.

#### 2.3.1 Stimuli

All participants underwent binocular visual field mapping using a luminance contrast retinotopic mapping (LCR). Figure 1A shows an example. Additionally, the glaucoma participants observed the LCR monocularly, and the healthy participants viewed the LCR binocularly with a simulated scotoma (LCR SS) superimposed (Figure 1B).

**Figure 1.**
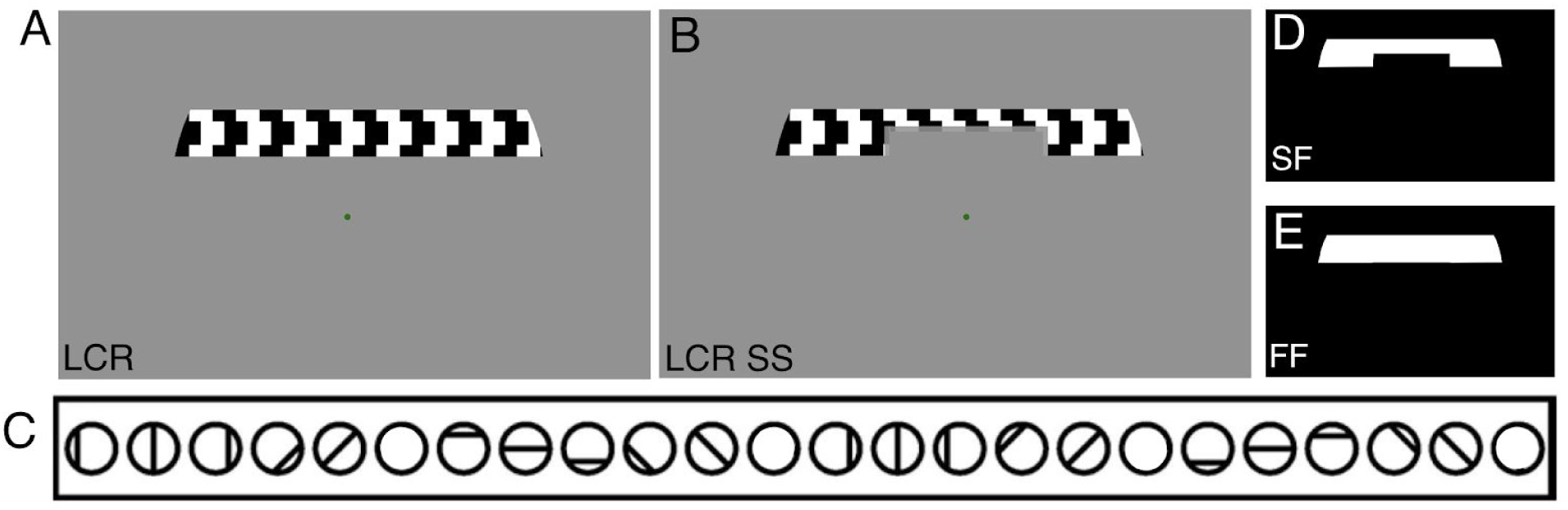
Example of the stimuli used to obtain pRF parameter estimates. **(A)** LCR stimulus. **(B)** LCR SS stimulus, this particular example depicts the simulated scotoma SS1. The colour of the fixation dot changed between red and green. **(C)** Scheme of the bar movements: four orientations in two opposing directions. **(D**,**E)** Visual stimuli models used during the pRF estimation: scotoma field (SF) and full field (FF) model.

##### 2.3.1.1 Luminance-contrast retinotopy (LCR)

LCR consists of a drifting bar aperture defined by high-contrast flickering texture ^8^. The bar aperture, i.e. alternating rows of high-contrast luminance checks drifting in opposite directions, moved in eight different directions: four bar orientations (horizontal, vertical, and the two diagonal orientations) and for each orientation two opposite drift directions. The bar moved across the screen in 16 equally spaced steps, each lasting 1 TR (repetition time, time between two MRI volume acquisitions). The bar contrast, width, and spatial frequency were 100%, 1.75 degree, and 0.5 cycles per degree, respectively. After each pass, during which the bar moved across the entire screen during 24 s, the bar moved across half of the screen for 12 s, followed by a blank full screen stimulus at mean luminance for 12 s as well, as shown in Figure 1.C.

##### 2.3.1.2 Luminance-contrast defined retinotopy with simulated scotomas (LCR SS)

LCR SS consisted of the LCR stimulus with a simulated scotoma superimposed. A total of six different scotomas were designed, one per participant. The different SS have irregular shapes and different sizes and were designed to mimic different clinical conditions. Figure 3A presents the six SS. SS1 is a central scotoma, as seen in age-related macular degeneration (AMD). SS2 and SS3: a central island and a nasal/arcuate scotoma, respectively, as seen in glaucoma. SS4-6 contain scotomas of different sizes and shapes, designed to further evaluate the method. The edges of the scotomas were smoothed using an exponential contrast mask (ECM), 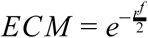, where *r* is the distance from the center of the scotoma and *f* is fixed at a value of 50.

**Figure 2.**
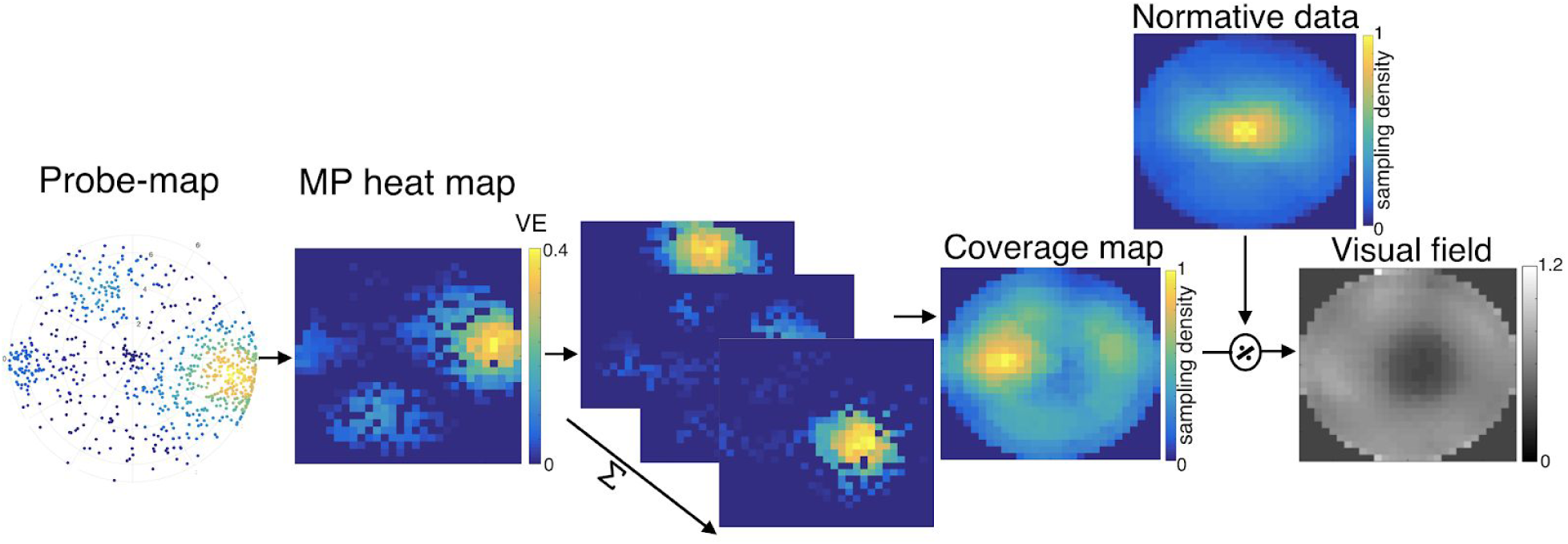
Pipeline of fMRI-based VF reconstruction using MP. First the probe map obtained with MP is converted into a heat map, a step done for every voxel within the cortical visual area of interest (e.g. V1). Next, these heat maps are averaged across all the voxels of that visual area, resulting in a mean VF coverage map. Finally, the reconstructed VF is obtained by dividing the individual normalized coverage map by the average normalized coverage map of all healthy participants, excluding the one in question (normative data).

**Figure 3.**
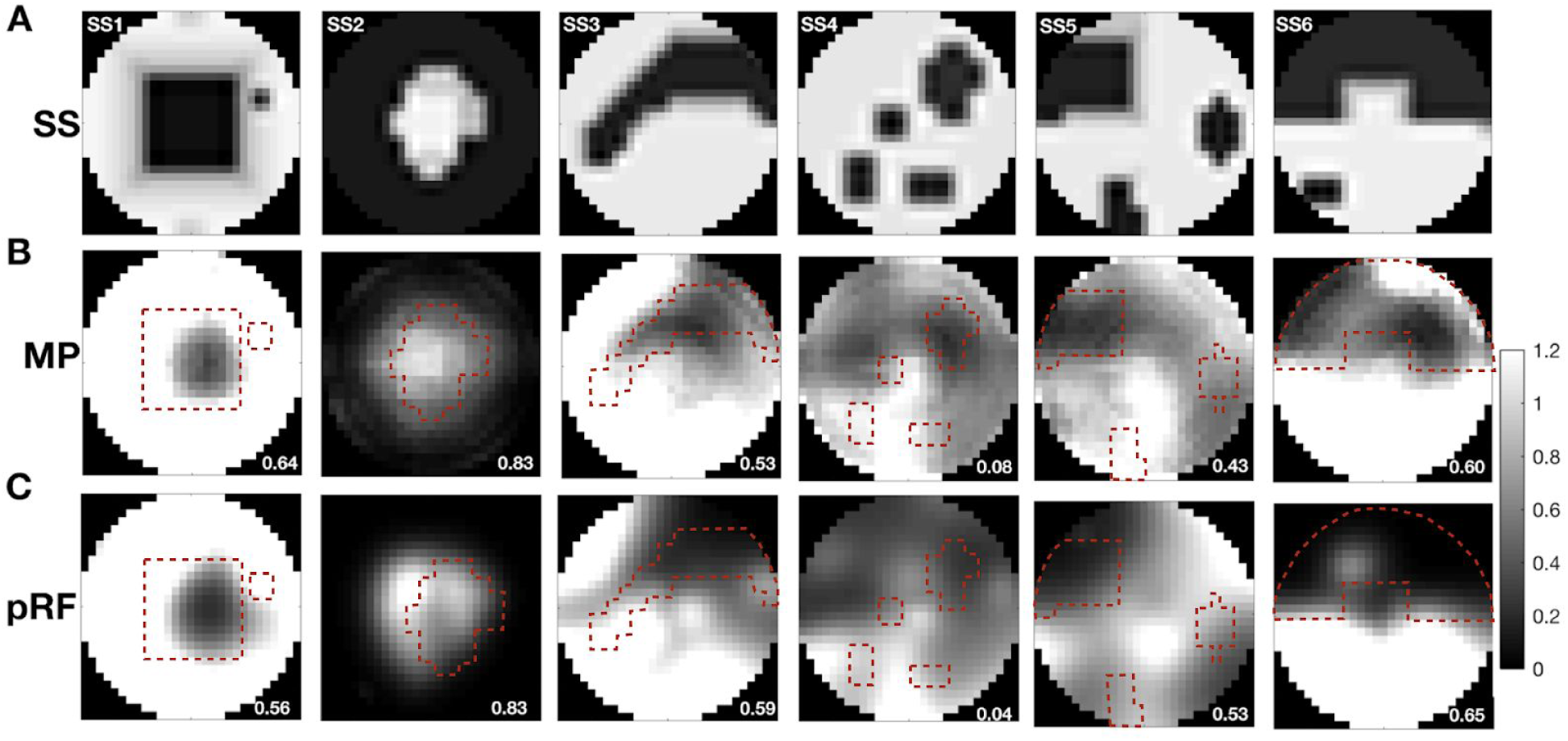
Visual field reconstructions based on retinotopic mapping fMRI data acquired in the presence of simulated scotoma. A: Representation of the different simulated scotomas. Dark regions correspond to low luminance contrast sensitivity. We used the following simulations: SS1: a central, square shaped scotoma of 10×10 deg; SS2: a peripheral scotoma with an irregularly shaped central island of approx. 8 deg diameter, SS3: a nasal/arcuate scotoma; SS4: four scotomas of different shapes: the smallest scotoma had a dimension of 1×1deg while the largest one measured 4×4 deg; SS5: three scotomas with different shapes: the smallest scotoma had a dimension of 1×1deg while the largest one was approximately the size of one quarterfield; SS6: two scotomas: a large one occupying the upper half of the visual field (with macular sparing) and a small one measuring 1×2 deg. B,C: visual field reconstructions based on V1 data using micro-probing (B) and conventional pRF mapping (C). The correlation between the simulated and reconstructed visual fields is shown in the bottom right corner of each reconstruction. The dashed red line corresponds to the edges of the SS overlaid with the VF reconstruction.

##### 2.3.1.3 Attentional task

During scanning, participants were required to perform a fixation task. The fixation task differed for healthy participants and participants with glaucoma. Healthy participants were instructed to press a button each time the fixation dot changed colour between green and red; participants with glaucoma were asked to press a button each time the fixation cross changed colour between black and yellow. The fixation cross extended towards the edges of the screen so that it could be used as a cue towards the screen’s center for the participants with a central scotoma (this was not needed in the healthy participants as we could project the fixation dot on the SS). The average performance - correct detection of the colour change of the fixation cross/dot - was above 78% for all conditions and all participants.

### 2.4 Magnetic resonance imaging

#### 2.4.1 Data acquisition and preprocessing

Scanning was carried out on a 3 Tesla Siemens Prisma MRI-scanner using a 64-channel receiving head coil. A T1-weighted scan (voxel size, 1mm^3^; matrix size, 256 × 256 x 256) covering the whole-brain was recorded to chart each participant’s cortical anatomy. Padding was used for a balance between comfort and reduction of head motion. The functional scans were collected using standard EPI sequence (TR, 1500 ms; TE, 30 ms; voxel size, 3mm^3^, flip angle 80; matrix size, 84 × 84 x 24). Slices were oriented to be approximately parallel to the calcarine sulcus. For all retinotopic scans (LCR, LCR monocular, and LCR SS, see section 2.2.4), a single run consisted of 136 functional images (total duration of 204 s). The (S)SPZ localizers consisted of 144 functional images (duration of 216 s).

The T1-weighted whole-brain anatomical images were reoriented in AC-PC space. The resulting anatomical image was automatically segmented using Freesurfer ^23^ and subsequently edited manually. The cortical surface was reconstructed at the gray/white matter boundary and rendered as a smoothed 3D mesh ^24^.

The functional scans were analysed in the mrVista software package for MATLAB (available at https://web.stanford.edu/group/vista/cgi-bin/wiki/index.php/MrVista). Head movement between and within functional scans were corrected (Nestares and Heeger, 2000). The functional scans were averaged and co-registered to the anatomical scan (Nestares and Heeger, 2000), and interpolated to a 1mm isotropic resolution. Drift correction was performed by detrending the BOLD time series with a discrete cosine transform filter with a cutoff frequency of 0.001Hz. In order to avoid possible saturation effects, the first 8 images were discarded.

#### 2.4.2 Visual field mapping

The pRF analysis was performed using both conventional population receptive field (pRF) mapping ^8^ and micro-probing ^25^. Using both the conventional and micro-probing models, for all the participants the functional responses to LCR and monocular LCR were analysed using a full field (FF) model, see Figure 1E. Additionally, the LCR SS condition was analysed using a model that used the SS stimulus mask as a prior (scotoma field; SF), see Figure 1D. The SF model was employed based on previous studies which proposed that the use of the SS as a prior into the pRF model mitigates methodological biases associated with the presence of scotomas ^11,19^. The prior knowledge about the simulated scotomas was applied in a similar manner to MP and pRF. In the fitting procedure of both pRF and MP, the overlap between the 2D Gaussian model- a probe (MP) and a candidate pRF model (pRF)- and the stimulus (binary mask of the stimulus over time *s*(*x, y, t*)), as described in the equation below:

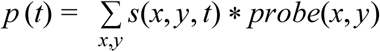

In practice, when we added prior knowledge about the simulated scotoma, the binary stimulus mask included the scotoma (as shown in figure 1D).

##### 2.4.2.1 Conventional pRF mapping

In the conventional method, a 2D-Gaussian model was fitted with parameters: center (x_0_, y_0_) and size (σ - width of the Gaussian) for each voxel. We used SPM’s canonical Haemodynamic Response Function (HRF) model. The conventional pRF estimation was performed using the mrVista (VISTASOFT) Matlab toolbox. The data was thresholded by retaining the pRF models that explained at least 15% of the variance.

##### 2.4.2.2 Micro-Probing

Micro-probing applies large numbers of “micro-probes”, 2D-Gaussians with a narrow standard deviation, to sample the entire stimulus space and create high-resolution probe maps. The number of micro-probes included, 10000, was calculated based on the trade off between achieving a good coverage of the visual field and the time to compute a probe map, Figure S1 and Table S2 ^26^. Like the conventional pRF approach, these micro-probes sample the aggregate response of neuronal subpopulations, but they do so at a much higher spatial resolution. Consequently, for each voxel, the micro-probing generates a probe map representing the density and variance explained (VE) for all the probes.

#### 2.4.3 ROI definition

The cortical borders of visual areas were derived based on phase reversal, obtained with the conventional pRF model using the LCR stimulus presented binocularly. Per participant, three visual areas (V1, V2, and V3) were manually delineated on the inflated cortical surface.

#### 2.4.4 Visual field coverage maps

The visual field representation at different levels of processing (low to high order areas) can be reconstructed by backprojecting the pRF responses obtained at the different ROIs onto the visual field. The visual field backprojection, also known as the coverage map, denotes the sampling density of the visual cortex. The reconstructed visual field maps were estimated using the conventional pRF mapping and micro-probing technique.

##### 2.4.4.1 Conventional pRF mapping

Using the conventional pRF, the coverage maps were estimated by including the pRF estimates whose VE was 0.15 or more. The percentage of voxels discarded by this threshold were 32% and 25% respectively for participants with glaucoma and normal vision participants. Per hemisphere and per visual areas, i.e, V1, V2, and V3, the pRF models were summed, weighted by their respective VE. By weighting the model by its VE, we take into account the responsiveness of the neuronal populations within a voxel and we reduce the effect of pRFs resulting from noise signals. Per visual area, a visual field coverage plot was then created for each individual participant by averaging the pRFs estimates for the left and right hemispheres. The described procedure is given by the following equation:

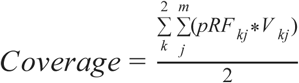

where *pRF* represents the clustering of all models from both hemispheres, *pRF*_*kj*_ is the model of the *j*-th voxel (*j*=1,…,*m*) in the *k*-th hemisphere (k=1,2) and *V* _*kj*_ its associated explained variance (* denotes convolution).

As a result of cortical magnification (resulting in a higher density of neurons in the areas representing the fovea than in those representing the periphery), the highest sampling density lies at the centre of the visual field. To correct for this tendency and to be able to straightforwardly recognize differences in pRF distribution between participants and as a consequence of (simulated) VFD, each participant’s coverage map was normalized using a normative dataset. In the case of the participants with normal vision, this dataset consisted of the average of the normalized coverage maps of all healthy participants (excluding the one in question) recorded for the unmasked condition. The normative dataset relative to the participants with glaucoma consisted of the average of the normalized coverage maps of nineteen aged-matched controls, the demographics of the normative cohort is shown on table1. Prior to this step it is necessary to normalize the coverage maps between 0 and 1, this allows us to take into account the different amount of voxels between each participant’s visual areas. Note that every visual area is defined specifically for every participant based on the retinotopic maps. This correction was applied in both methods (pRF and MP).

##### 2.4.4.2 Micro probing

For each voxel, the micro-probing generates a probe map representing the density and VE for all the probes. These probe maps were converted into single voxel coverage maps - heat maps (2 dimensional histograms of a 30 × 30 bin grid weighted by its bin variance explained). At the level of V1, the visual field can be reconstructed by summing the V1 heat maps across all V1 voxels.

The final average coverage map included all V1 voxels for which the heat map had at least one location (bin) for which the VE was higher than 0.15. The percentage of voxel discarded by this threshold was 33% and 28% respectively for participants with glaucoma and normal vision participants. Note that the discarded voxels correspond to the ones where the MP did not converge, which is most likely due to noisy measured signals. This threshold is equivalent to the one applied in standard analysis of the pRF modelling (section 2.4.4.1, ^27,28^). Figure 2 represents a scheme of the steps underlying the VF reconstruction using micro-probing.

### 2.5 Statistical analysis

As in previous work ^8,14,29^, data was thresholded by retaining the pRF models that explained at least 0.15 of the variance in the BOLD response and that had an eccentricity in the range of 0-7 degrees (this is a necessary step to ensure the estimated pRF is located within the stimulated field of view), for all conditions (i.e., LCR and LCR SS). The correlations between the contrast sensitivity loss and the sampling density obtained with MP and conventional pRF mapping were calculated using a linear mixed effects model with a slope and intercept per participant as a random effect.

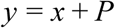

where *y* is visual sampling, *x* is contrast sensitivity loss and P the random effect of each participant. The contrast sensitivity loss of a visual field area, i.e, a quadrant of 7×7 deg adjacent to fixation, was calculated from the total deviation values of the SAP test locations. This was done by up-sampling the SAP resolution to 1 deg^2^ and subsequently allotting to each 1 deg^2^ the sensitivity of the nearest SAP test location, using the locations at (±3,±3), (±3,±9), (±9,±3), and (±9,±9) deg (with the origin at fixation).

## 3. Results

### 3.1 fMRI-based visual field reconstruction of heterogeneous simulated visual field defects

Figure 3 shows how the VF reconstructions based on MP (panel B) and pRF mapping (panel C) correspond to the SS (panel A). For each scotoma, the pearson correlation coefficients between the SS and reconstructed VF are shown in Table 2 and also on the reconstructed images of Figure 3. For VF reconstructions based on V1 activity, the median correlation between the SS and reconstructed VF is 0.57 and 0.58, for MP and pRF respectively. Large scotoma (> 3 deg) are better reconstructed than smaller ones. SS2, a simulation of tunnel vision with a large peripheral scotoma, has correlations of 0.83 for both MP and pRF, while SS4, with small scotomas, shows correlations of only 0.08 and 0.04 for MP and pRF, respectively. Both methods primarily detect scotomas with a diameter larger than 3 deg (see e.g. SS4).

**Table 1.** Demographics of the participants with glaucoma and of the normative cohort. Average and standard deviation of age, percentage of female, intraocular pressure (IOP) for the right and left eye (average over three measurements), peripapillary retinal nerve fiber layer (pRNFL) thickness and the VF mean deviation (VFMD) measured with SAP for the right and left eye. Note that participants with glaucoma were receiving treatment.

| Measure | Participants with Glaucoma |  | Normative cohort |  |
| --- | --- | --- | --- | --- |
|  | Average (n=19) | Standard Deviation (n=19) | Average (n=19) | Standard Deviation (n=19) |
| Age (y) | 70 | 8.8 | 68 | 7.3 |
| Gender (F%) | 52 | - | 42 | - |
| IOP (R/L; mmHg) | 13.4/13.4 | 2.6/3.9 | 13.1/13.5 | 2.8/3.2 |
| pRNFL thickness (R/L; $\mu\text{m}$) | 72.7/ 68.0 | 11.7/9.6 | 96.84/97.2 | 9.7/10.9 |
| VFMD (R/L; dB) | -7.4/-9.1 | 8.1/8.3 | -0.4/-0.69 | 1.4/3.4 |

**Table 2.** Correlations between simulated and reconstructed VF for the visual areas V1, V2, V3 and aggregating the visual areas using in the pRF analysis the full field model (FF) and a model that uses the SS stimulus mask as prior (SF). A indicates reconstructed VF on aggregated V1-V3 data.

|  | Reconstruction method, analysis model and visual area |  |  |  |  |  |  |  |  |  |  |  |  |  |
| --- | --- | --- | --- | --- | --- | --- | --- | --- | --- | --- | --- | --- | --- | --- |
|  | Micro-probing |  |  |  |  |  |  | Conventional pRF |  |  |  |  |  |  |
|  | FF |  |  |  | SF |  |  | FF |  |  |  | SF |  |  |
| Simulation | V1 | V2 | V3 | A | V1 | V2 | V3 | V1 | V2 | V3 | A | V1 | V2 | V3 |
| SS1 | 0.64 | 0.51 | 0.34 | 0.72 | 0.80 | 0.75 | 0.70 | 0.56 | 0.34 | 0.27 | 0.56 | 0.57 | -0.20 | -0.03 |
| SS2 | 0.83 | 0.57 | 0.73 | 0.74 | 0.94 | 0.84 | 0.90 | 0.83 | 0.65 | 0.78 | 0.79 | 0.20 | 0.25 | 0.59 |
| SS3 | 0.53 | 0.41 | 0.35 | 0.60 | 0.67 | 0.65 | 0.63 | 0.59 | 0.45 | 0.30 | 0.56 | 0.48 | 0.12 | -0.30 |
| SS4 | 0.08 | 0.10 | -0.07 | 0.04 | 0.52 | 0.57 | 0.57 | 0.04 | -0.19 | -0.02 | -0.23 | -0.15 | -0.43 | -0.60 |
| SS5 | 0.43 | 0.19 | 0.17 | 0.29 | 0.76 | 0.65 | 0.61 | 0.53 | 0.16 | 0.39 | 0.49 | 0.24 | -0.15 | 0.13 |
| SS6 | 0.60 | 0.67 | 0.53 | 0.72 | 0.84 | 0.86 | 0.69 | 0.65 | 0.69 | 0.73 | 0.78 | 0.64 | 0.48 | 0.63 |
| Median | 0.57 | 0.46 | 0.35 | 0.66 | 0.78 | 0.70 | 0.66 | 0.58 | 0.40 | 0.35 | 0.56 | 0.36 | -0.02 | 0.05 |

To infer the effect of between-subject variability, we calculated the correlation between each participant’s reconstructed VF without simulations and the average reconstructed VF of the remaining participants for the visual areas V1, V2, and V3 using MP (median correlation across visual areas: 0.91) and pRF (median correlation across visual areas: 0.81). Conventional pRF mapping (SD across visual areas: 0.06) resulted in a larger between-subject variability than MP (SD across visual areas: 0.15). Table 3 presents the results for each participant obtained using MP and pRF.

**Table 3.**
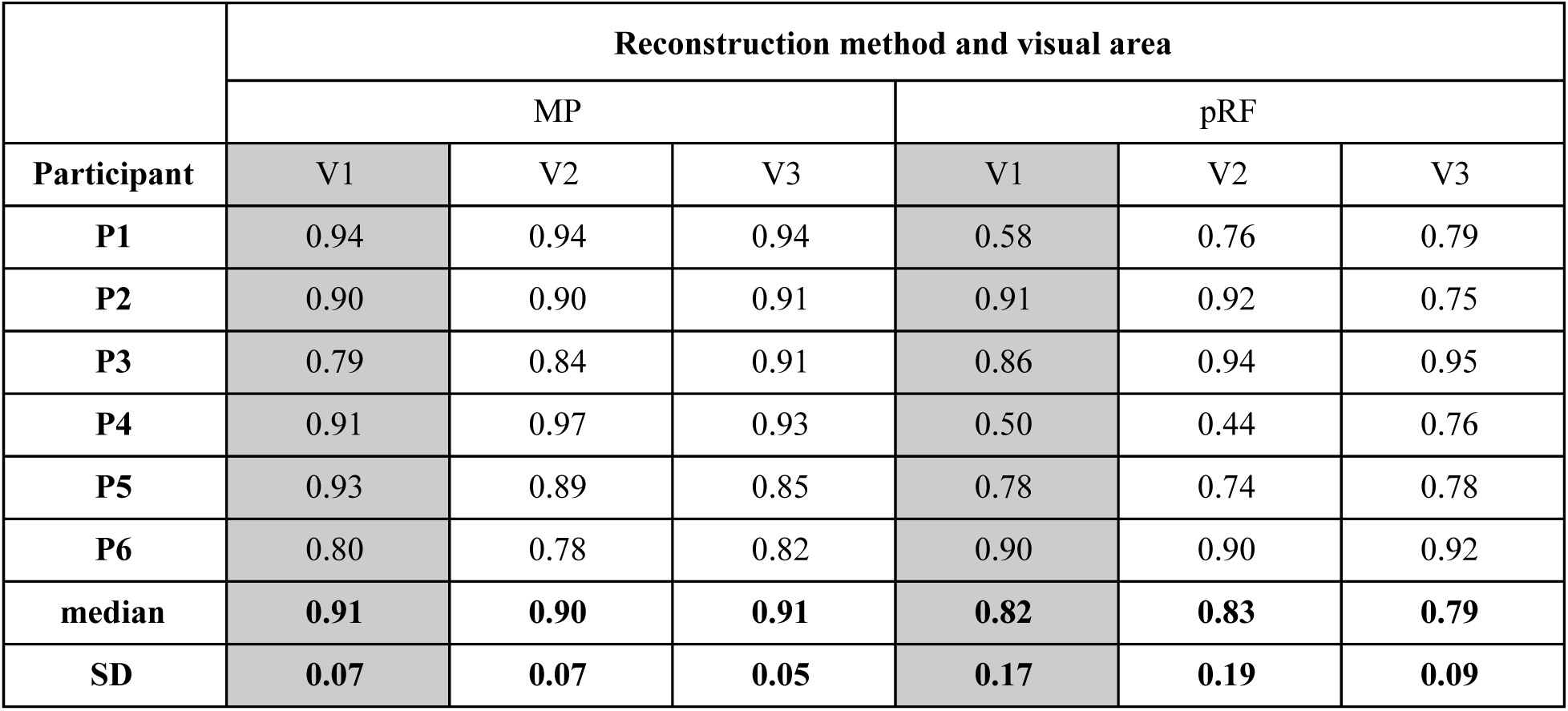
Correlation between each participant’s reconstructed VF without SS and the reconstructed VF of the remaining participants for the visual areas V1,V2 and V3 using MP and the conventional pRF. The average and the standard deviation (SD) across participants are presented in the bottom rows. Note that, a priori, for both models MP and pRF the correlation between the reconstructed visual field and the SS mask is expected to be smaller than the correlation between the visual field reconstructions.

### 3.2 Shrinkage of visual field defects across the visual hierarchy

Figure 4 shows the reconstructed visual field based on the pRF estimates obtained with the conventional pRF approach for the visual areas V1, V2 and V3. Figure 5 shows the same but for MP. It is clear that the VFD becomes smaller along the visual hierarchy, which is translated in a decrease of the correlation values between the scotoma mask and the reconstructed visual field. This effect is less pronounced for the MP than for the pRF-based approach. Using both techniques, the differences between SF and FF models are accentuated for the visual areas V2 and V3 compared to V1. For MP, this most likely results from the fact that using the FF model, with the increase in the visual hierarchy the reconstructed VFD become smaller and therefore the correlation between simulated and reconstructed VF is lower, whereas using the SF the VFD does not shrink with visual hierarchy resulting in a higher correlation. For the conventional pRF the differences are likely to result from the biases associated in the prior models of the scotoma.

**Figure 4.**
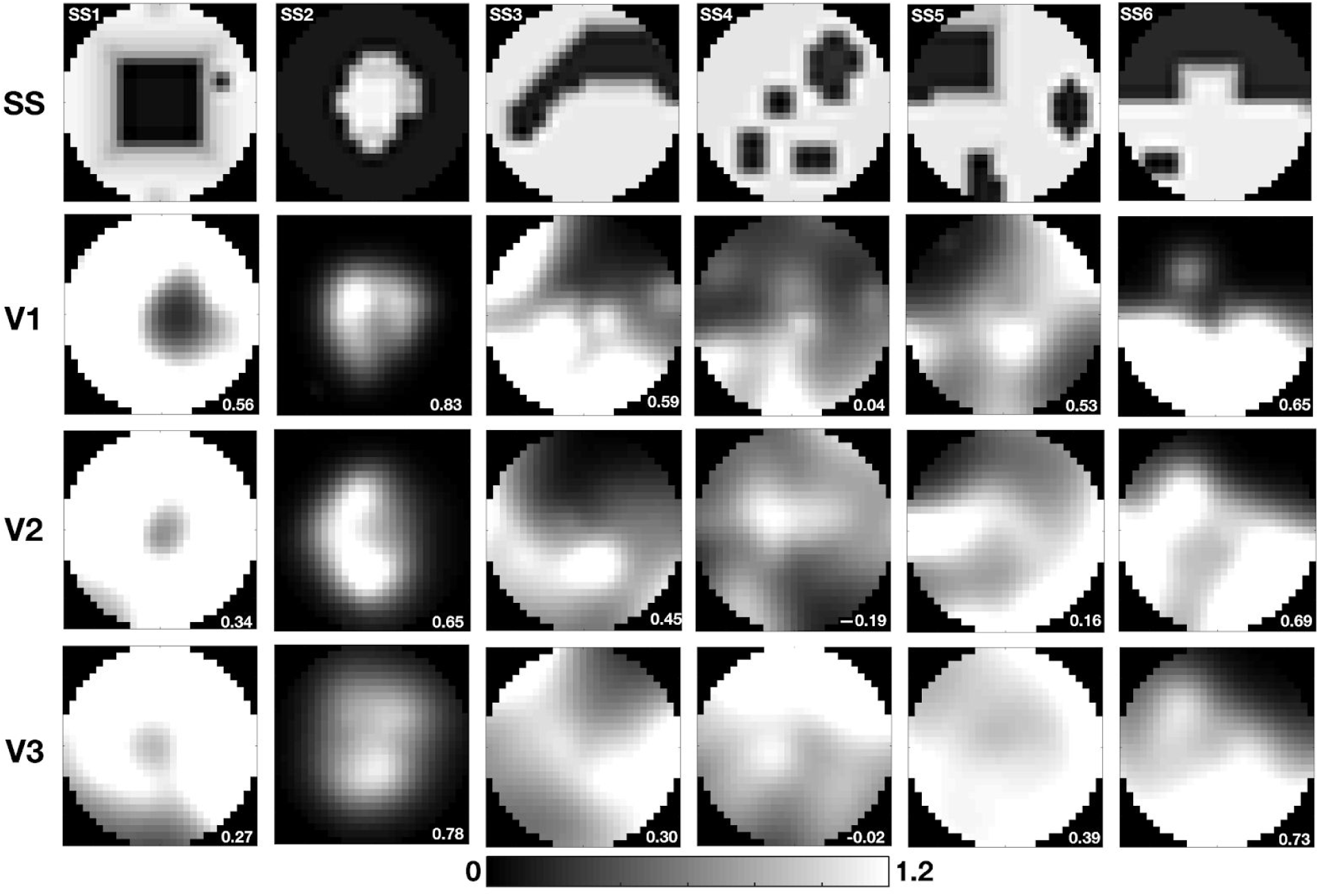
Visual field reconstruction in the presence of simulated visual field defects at different levels of the visual cortical hierarchy. Visual field reconstruction based on conventional pRF mapping for V1, V2, and V3. The correlation coefficients (also presented in Table 2) between the reconstructed visual field and the simulated scotoma used are provided in the bottom right corner of the images.

**Figure 5.**
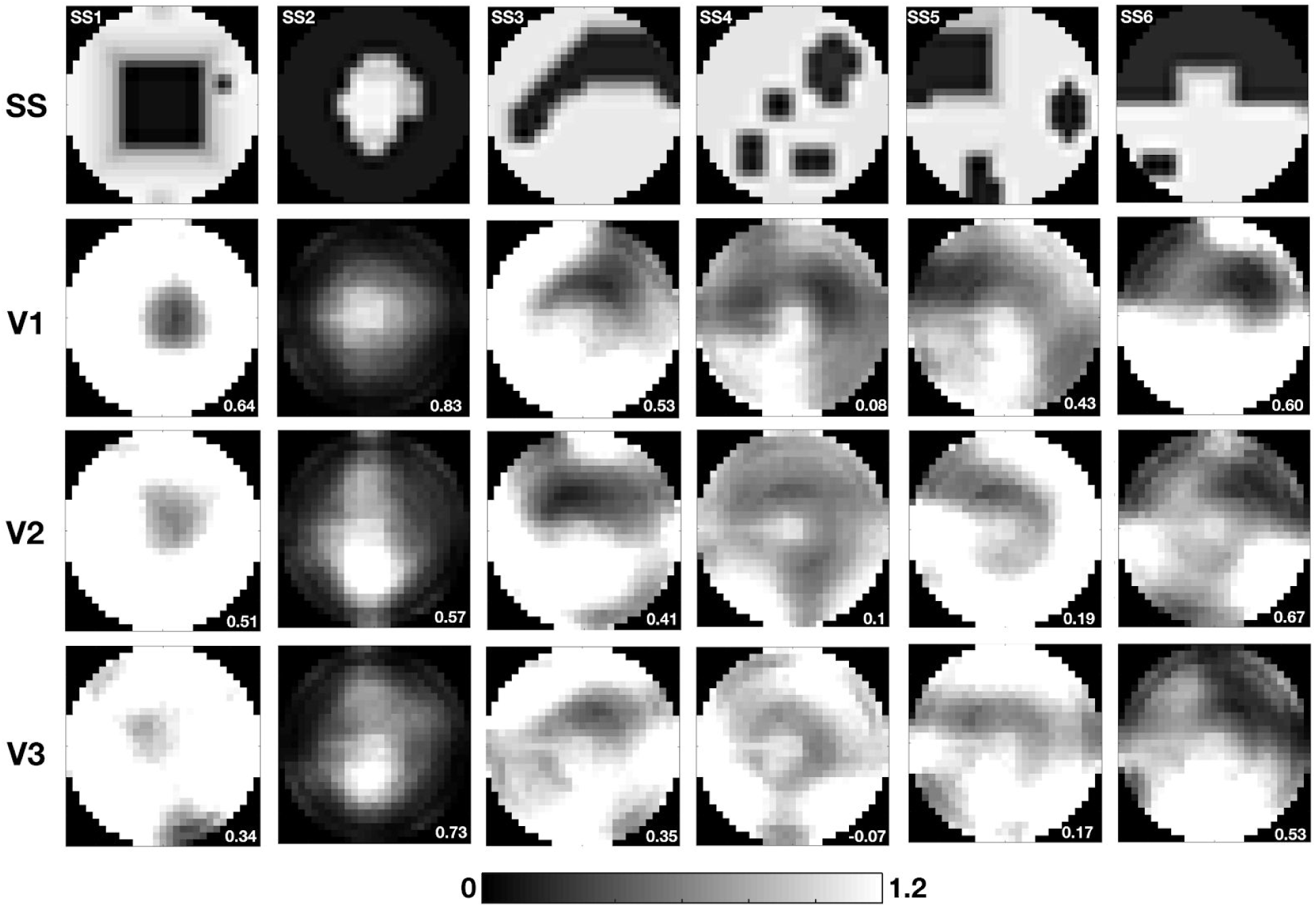
Visual field backprojection using simulated visual field defects across the cortical hierarchy using MP. Visual field reconstruction based on MP for V1, V2, and V3. The correlation coefficients (also presented in Table 2) between the reconstructed visual field and simulated scotoma used are presented in the bottom right corner of the images.

In addition, we reconstructed the visual field combining the data obtained for V1, V2, and V3. The results are shown in Figure 6. Compared to the correlations obtained exclusively based on V1, there is no clear benefit of reconstructructing the VF based on the aggregated data across areas (MP t(5,5)=0.155, p=0.88; pRF t(5,5)=0.26, p=0.8). The correlation between the simulation and reconstructed visual field across areas is slightly higher for MP (0.66) than for the pRF (0.56), to be compared to 0.57 (MP) and 0.58 (pRF) for V1 alone (Table 2). However there are no significant differences between the correlation of the reconstructed VF obtained with MP and pRF (V1 level t(5,5)=-0.28, p=0.79, combined V1, V2,V3 t(5,5)=0.16, p=0.87).

**Figure 6.**
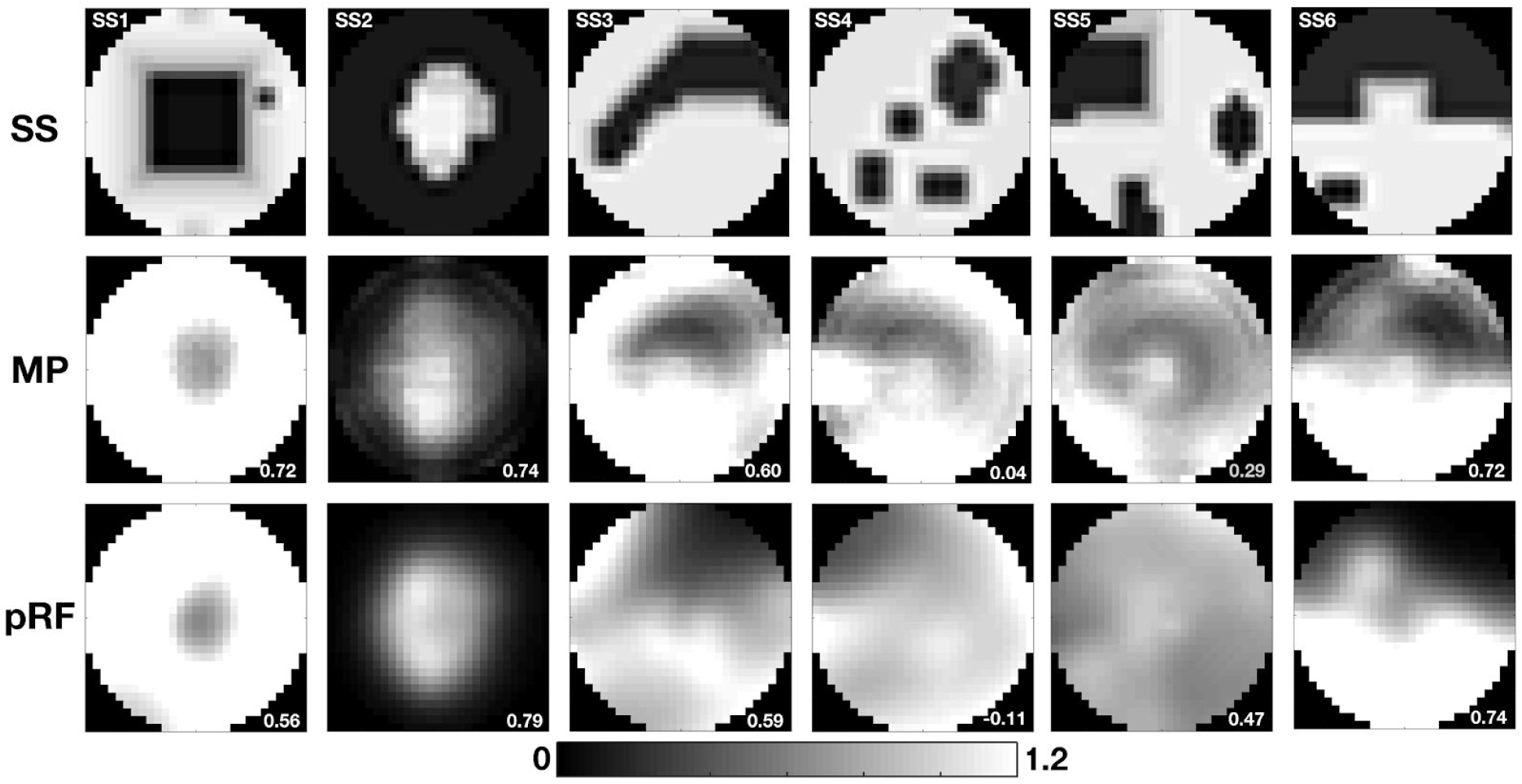
Visual field reconstruction of simulated scotomas based on data aggregated across V1, V2, and V3. Visual field reconstruction based on MP and conventional pRF (both using FF model) obtained by averaging the V1, V2 and V3 visual field maps. The correlation coefficients (also presented in Table 2) between the reconstructed visual field and the simulated scotoma are presented in the bottom right corner of the images.

### 3.3 Effect of providing prior information on the location of SS during pRF estimation

A major difference between MP and conventional pRF models relates to how the VF is sampled. In pRF procedure the 2D Gaussian can be located in the scotoma and extend towards the spared VF, this will result in a partial stimulation of the pRF, which can be interpreted as sampling from within the scotoma region. In contrast, in MP the probes are very small, strongly reducing the probability that they are located inside the scotoma and still be partially stimulated. To understand if the use of MP and the incorporation of prior knowledge about the SS location into the pRF model reduces the biases in pRF estimated properties, we applied two different models. The full field (FF) model did not include the simulation, while the scotoma field (SF) model did. However, it is important to note that for VF reconstruction the adding prior knowledge about the location of the scotomas is not feasible in clinical conditions. Figure 7 shows the reconstructed visual field based on V1 data taking into account the scotoma definition during the pRF modelling via MP and pRF. Panel A shows that when using MP in combination with the SF model, there is little to no sampling of the scotomatic region and, consequently, the reconstructed VFs better correlate with the SS (median correlation: 0.78) compared to the FF model (median correlation: 0.57), (t(5,5)=-4.43, p=0.007). In contrast, using the pRF there is residual sampling within the scotomatic region, most likely resulting from large pRFs located outside of the scotoma (panel B). Notably, for pRF the quality of the reconstruction is slightly lower using the SF model (median correlation: 0.36) compared to the FF model (median correlation: 0.53) (t(5,5)=2.10, p=0.09).

**Figure 7.**
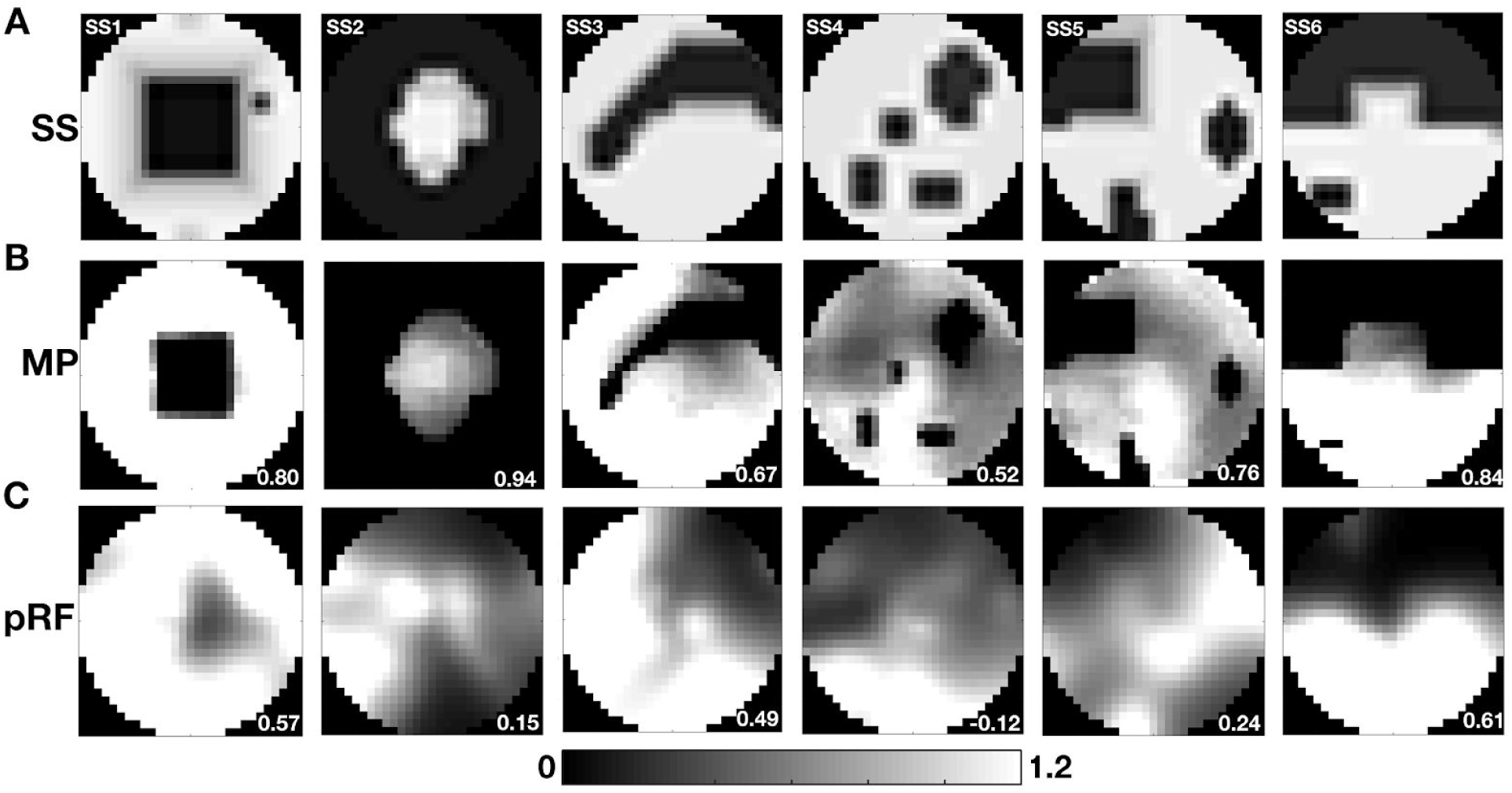
Visual field reconstruction when including the scotoma into the analysis model (SF). A: Simulated VFD. The two rows below show visual field reconstructions based on V1 data using either MP (A) or pRF mapping (B). The correlation between the simulated and reconstructed visual field are shown in the bottom right corner of each reconstruction.

Importantly, the comparison of the spared VF (region of the VF without SS) between SF and FF models for MP and pRF approaches showed a higher correlation for MP (0.99) than for the conventional pRF (0.89), (t(5,5) = 3.74, p = 0.0139). The correlation coefficients obtained for each participant are shown on Table S2. This indicates that MP is more robust to modelling biases than the pRF approach.

### 3.4 fMRI-based visual field reconstruction enables the detection of natural scotomas

To understand if pRF based techniques can be used to detect natural scotomas, we applied MP and pRF to the monocular retinotopic data of a group of glaucoma participants. We qualitatively compared the visual field reconstructions with their perimetric measurements, obtained using SAP. Figure 8 shows the fMRI-based visual field reconstruction next to the SAP-based VF and the macular ganglion cell complex (GCC) thickness for 19 participants with glaucoma. The gray scale SAP VF map represents the contrast sensitivity values measured in decibels (dB) at multiple locations of the VF. Normal contrast sensitivity values are around 30 dBs, thus lower sensitivities are indicated by darker areas (black corresponds to 0 dB) and higher sensitivities are represented with a lighter tone (<30 dBs are represented in white). The OCT maps are centered in the fovea and represent the macular thickness values in microns compared to a normative dataset. The thinnest 1% of measurements fall in the red area which is considered outside normal limits. Visually, one can appreciate that MP and pRF capture coarsely the same patterns of organization and the high between-participant variability between the SAP VF and the fMRI-based VF. While for some participants these are very similar (participant G12) for others there is a clear dissociation between the two techniques (participant G14). Note that the SAP field of view shown in Figure 8 is much larger than the one of the fMRI-based VF reconstruction (7 deg), with the red circle denoting the corresponding field of view.

**Figure 8.**
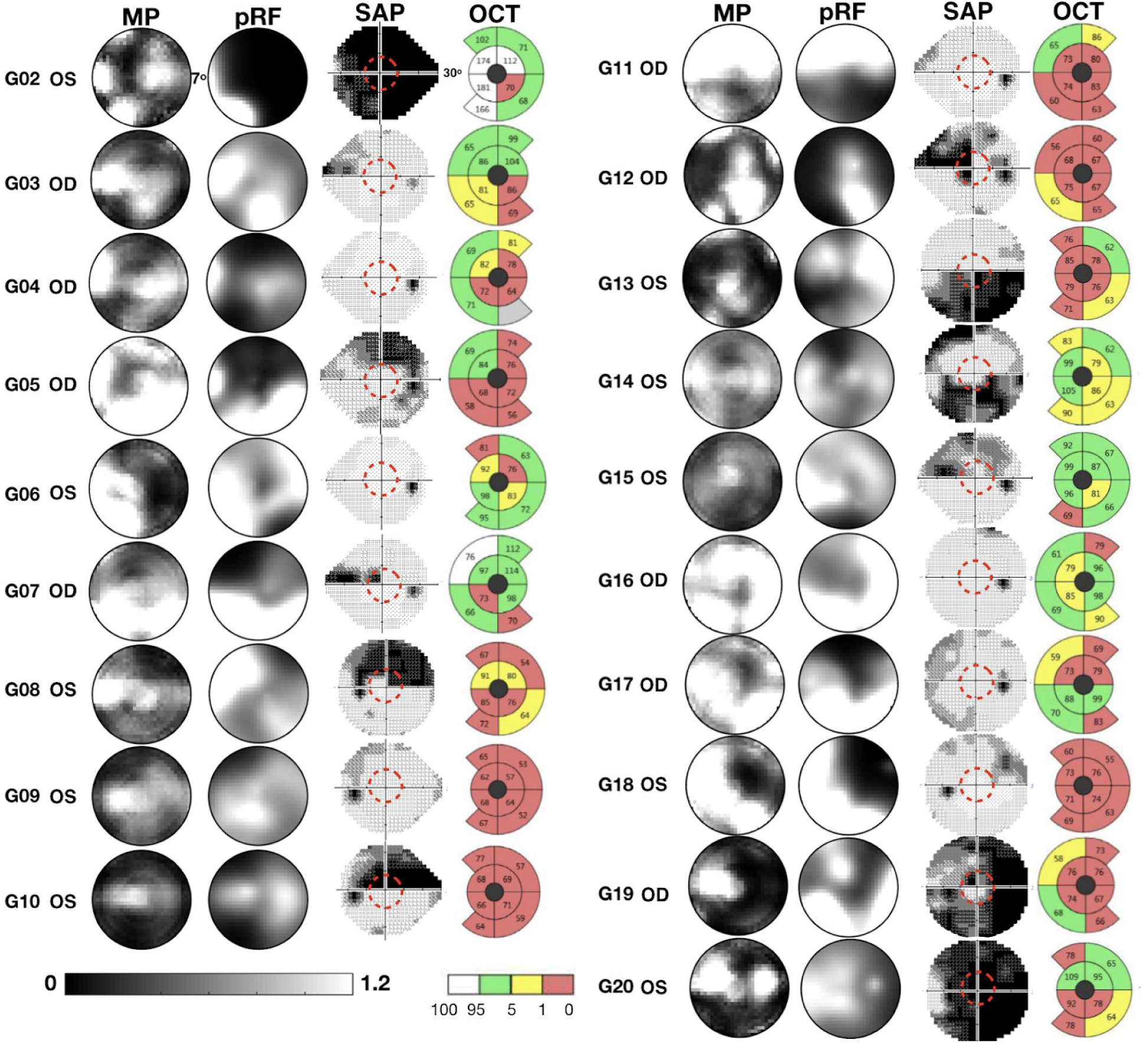
Reconstructed VF using MP and conventional pRF, SAP, and the OCT-derived GCC thickness for 19 glaucoma participants, for the most affected eye (based on MD - an overall measure that indicates how much a participant deviates from an age-matched normative data set). The red dashed circle denotes the field of view of the fMRI-based approaches (7 deg). The OCT image covers about 20 deg. White, green, yellow and red colours of the OCT maps correspond to the thickest 5%, 90%, thinnest 5% and thinnest 1% of measurements. A shaded gray area corresponds to a disk area outside the central 90% of normal range.

#### 3.4.1 Micro-probing better predicts the contrast sensitivity than conventional pRF mapping

We quantified the degree of similarity by calculating the correlation between the sampling density of individual VF quadrants measured with MP and pRF mapping and contrast sensitivity loss scores obtained with SAP. Figure 9 shows that the sampling density obtained with both methods correlates significantly with SAP-based contrast sensitivity, with MP providing a higher correlation (MP: r^2^=0.44, p=0.0002; pRF: r^2^=0.32, p=0.003). Model comparison using Bayesian Information Criteria (BIC) showed that MP (BIC=592) is significantly better than the conventional pRF(BIC=600) in predicting contrast sensitivity.

**Figure 9.**
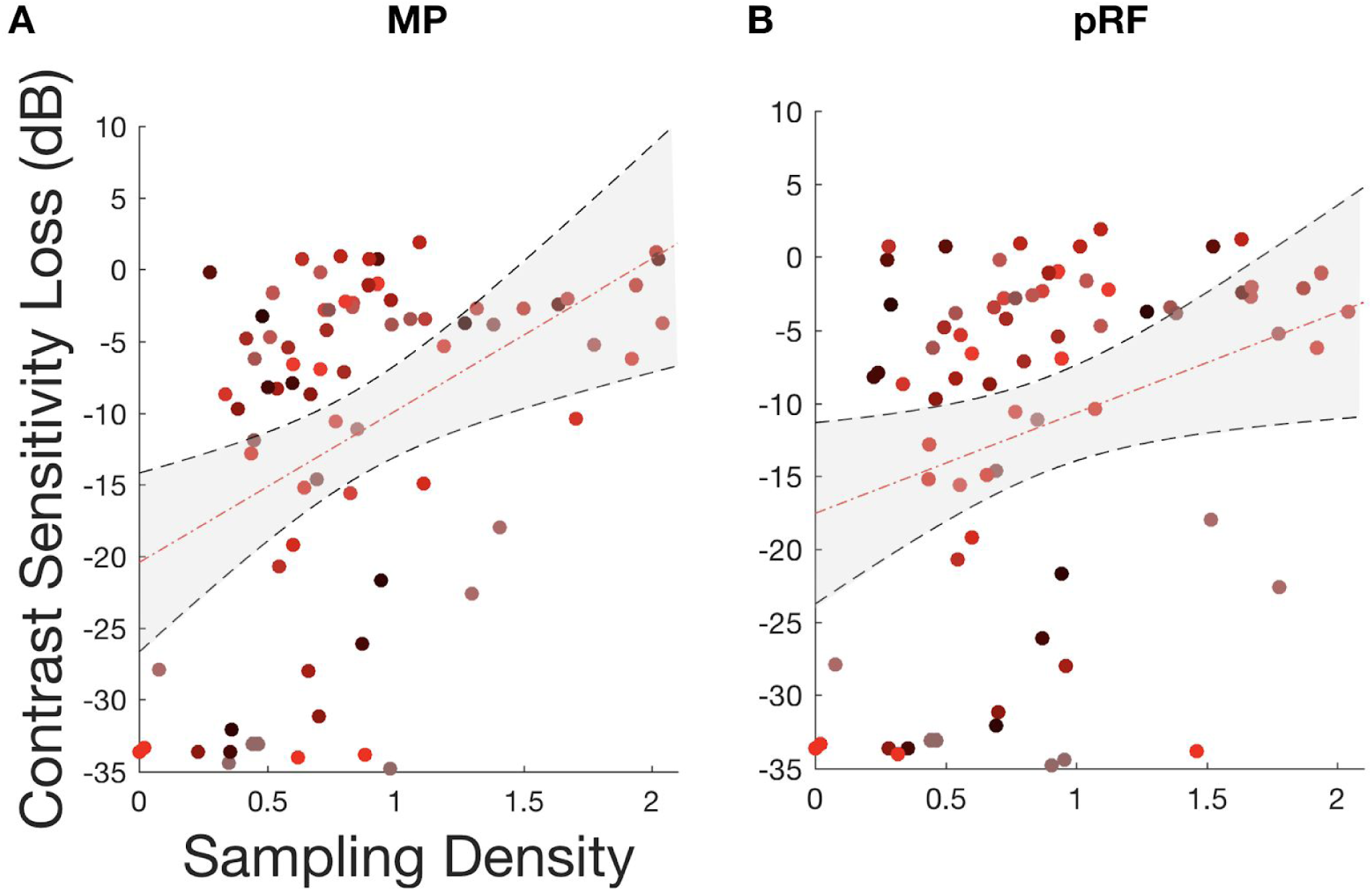
Correlation of sampling density measured with fMRI-based techniques and contrast sensitivity loss obtained with SAP. Panels A and B show the correlation of the sampling density of participants with glaucoma obtained from individual quadrants with contrast sensitivity loss, for MP (A) and pRF (B) techniques, respectively. Each data point is from a separate quadrant of an individual participant with glaucoma. Each color represents the datapoint of each participant. The dashed red line represents the linear fit to the data and the shaded region represents the 95% confidence interval of the fitted parameters. Note that the correlations were obtained using the same visual field area for the fMRI-based visual sampling and the SAP-based contrast sensitivity (a 7×7 deg quadrant adjacent to fixation).

## 4. Discussion

Our main finding is that fMRI in combination with the MP and pRF analyses makes it possible to reconstruct the VF and to detect even fairly small defects (> 3 deg) therein. Importantly, this could be achieved in single participants, based on 12 minutes of effective fMRI scan time. When applied to participants with glaucoma, the fMRI-based VF reconstruction detected scotomas with a correspondence to SAP that was decent. Moreover, we found a moderately good correlation between fMRI-based sampling density and SAP-based contrast sensitivity loss, this correlation was better using our new approach than using the conventional pRF analysis. We will discuss the possibilities and limitations of the fMRI-based approach of VF reconstruction.

### 4.1 fMRI-based visual field reconstruction enables VFD detection and the accurate assessment of the visual field

Using quantitative retinotopic mapping (conventional pRF and MP), we reconstructed the visual field of six participants with heterogeneous simulated scotomas at different levels of the visual processing hierarchy. Based on V1 data, we demonstrated the accuracy of the technique to reconstruct the visual field and detect simulated scotomas with diverse shapes and sizes.

However, the technique cannot assess the presence of small VFD (< 3 deg diameter). Our findings corroborate previous fMRI studies using simulated central scotomas and high spatial resolution protocols ^15,18^. We found that the reconstructed VFDs tend to become smaller when based on data of visual areas higher up along cortical hierarchy. This is most likely a consequence of the increased size of pRFs from early to higher order visual areas ^30^. The decrease in VFDs is more pronounced using conventional pRF than MP, this is most likely related with the fact that MP is more conservative estimating the size of pRFs ^26^. Additionally, we showed that, in the context of retinal damage, there is no benefit of combining the information from different visual areas. However, information about the visual representations across cortical hierarchy might be useful to access the efficacy of training, when applied as rehabilitation strategy in visual impairment caused by brain injuries (e.g., hemianopia).

### 4.2 To include or not to include the simulated scotoma into the model as a prior?

Baseler et al. found changes in pRF properties in healthy participants similar to those present in participants with macular degeneration ^14^. This indicates that a mere change in visual stimulation can give rise to changes in measured pRF properties, invalidating a direct interpretation of such changes in terms of plasticity ^12,31^). Consequently, it is critical to disentangle changes in pRF properties driven exclusively by changed visual input and arising from plasticity of the visual pathway following damage to the visual system ^11,12,14^. Accurate SS are thus a critical tool to help establish the presence of plasticity. Our results show that this is possible to achieve, provided the SS do not become too small.

The presence of scotomas results in partial stimulation of pRFs and in modulation of the pRF responsive to the scotoma location via feedback signals from higher order visual areas, which may lead to biases in the pRF estimates, i.e enlargement of the pRFs and shifts out of the scotoma zone ^11,12^. Previous studies proposed that the use of the SS as a prior into the pRF model mitigates these methodological biases. We investigated how our MP- and pRF-based VF reconstructions are influenced by incorporating prior knowledge on the presence of scotomas.

When using the FF model, MP and pRF perform similarly. However, when using the SF model, the difference between MP and pRF is more pronounced. Using MP in combination with an SF model ensures that no information is sampled from the scotomatic zone. Consequently, it can be an alternative approach to pRF linear models that enforce an absence of sampling of the SS projection zone, e.g., the Inverse model and the Linear Artificial Scotoma model ^11,19^. In contrast, reconstruction of the VF based on the pRF using an SF model results in sampling of the scotomatic region. Most likely, this sampling occurs via large pRFs located outside of the scotomatic zone. Moreover, based on the correlation between the stimulus mask and VF representations obtained with the FF and SF model based on the pRF model we found that the VF reconstruction based on the FF model is actually better than those obtained with the SF model. We interpret this as evidence that the conventional pRF model, when applying an FF model, compared to an SF model, actually reduces biases. This contrasts with the conclusions of previous studies which suggested that biases at the scotoma border can be ameliorated by incorporating the SS as a prior to the model when estimating the pRFs ^11,19^. These biases may result from a delay in response of voxels located near the scotoma edges caused by BOLD spreading or feedback from higher cortical areas, resulting in pRFs with more eccentric locations and larger sizes being assigned to those voxels ^11,12^. Due to its use of small sized probes that are more robust to partial stimulation effects, MP results in a less biased estimation of the preserved VF.

### 4.3 FMRI-based visual field reconstruction provides additional information to standard perimetry in the clinical assessment of a VFD

Applied to natural visual field defects of patients with glaucoma, overall the fMRI-based visual field reconstruction techniques showed a decent correlation with the SAP contrast sensitivity loss, and found the larger VFDs also detected by SAP. In particular, our new approach (MP) predicted the SAP-determined contrast sensitivity loss better than the conventional pRF approach did. However, for some of the participants with glaucoma, there was a dissociation between the reconstructed visual field using fMRI and the measured visual field using SAP. Most likely this dissociation is associated with differences in methodology. SAP uses spots of light close to threshold as stimuli whereas the present fmri-based approaches use a high-contrast (supra-threshold) moving luminance contrast bar. This has various potential implications. First, the bar may enable extrapolation (i.e. predictive masking) whereas the spots will not. The fMRI approach may thus underestimate the actual size of the scotoma. Second, it could also imply that the bar stimuli inform on active cortical tissue that is not detected by SAP. Third, fMRI-based approaches may be able to detect the macular vulnerability zone - an area of the superior paracentral VF often affected in early glaucoma - that cannot be detected with SAP due to the coarse grid (6×6 deg) sampling ^32^. The presence of OCT abnormalities within the macular area in many patients with a normal central visual field (Fig. 8) supports this hypothesis.

The combination of fMRI and neurocomputational models: 1) results in an objective measure of the visual field which complements current ophthalmic evaluations and 2) reveals potentially important characteristics of visual system functioning that cannot be assessed via or inferred from the standard ophthalmic examinations. These characteristics may become important when evaluating the impact of visual restoration and rehabilitation therapy on visual processing beyond the retina ^33^. Importantly, the reconstructed VF were obtained during only 12 minutes of data acquisition (this corresponds to effective visual field mapping time, it excludes time required to acquire the anatomical scan and the preparation of the participant to perform an MRI scan). Being time effective is a crucial aspect for the feasibility of applying this approach in clinical practice. Moreover, the acquisition time can be further reduced by using higher field strengths.

### 4.4 Limitations

In this study we used a 24-2 and 30-2 grid sampling HFA tests for the following reasons :1) the VF of glaucoma patients was checked every 6 months in the hospital using the 24-2 and 30-2 protocols, given the variability associated with these test, using the same protocol allowed to compared with previous exams and verify if the measure that we obtained were reliable, 2) we wanted to investigate if peripheral visual field defects could affect the visual sampling of the central visual field and 3) this study was part of a battery of tests also applied to aged matched controls, we deemed necessary to apply the same protocol to glaucoma and aged-matched control participants. A 24-2 or 30-2 grid is necessary to screen the aged-matched controls for small peripheral visual field defects. However, the relatively coarse sampling of the SAP luminance sensitivity (one measurement per 36 deg^2^) combined with the limited size of the screen inside the scanner (7 deg eccentricity) compromised the assessment of the accuracy of the visual field reconstruction techniques at an individual level; there were only few data points to establish the similarity between SAP and visual field reconstruction. This required upsampling the resolution of the SAP, which might lead to the attribution of an interpolated rather than an actual sensitivity at certain locations of the VF, in particular for positions at the edge of the 36 deg^2^ measurements of SAP. One way to mitigate this in future experiments is to use a wider MRI visual field of view and a finer grid SAP protocol, i.e., a 10-2 grid, or micro perimetry ^17^. Early stage glaucoma is characterized by loss of VF primarily in the periphery. Therefore, a limited screen size inside the scanner will limit the ability to detect peripheral VFD and thus the use of fmri-based VF reconstruction techniques in such cases. This limitation can be surpassed using MRI compatible goggles or adapted systems for wide-field visual field stimuli which can result in a stimulation upto (60 deg eccentricity)^34^.

Each participant only viewed one simulation. Therefore, differences in the accuracy of the reconstruction of the different SS may be associated with individual participant differences. However, all the participants included in the study with SS were experienced with retinotopic mapping. Moreover, the fact that the reconstructed visual field maps without SS were highly correlated between participants (MP: 0.90 and pRF: 0.81) suggests that, although inter-participant variability might play a role, overall it cannot explain the large differences in reconstruction accuracy of the different SS. Moreover, we interpret the overall agreement between each individual simulated SS and reconstructed VF as an indication that our approaches can accurately detect scotomas at the participant level.

Although an AS is considered a good model of visual field defects, there are various fundamental differences with natural scotoma that may limit the translation of our present findings. First, natural scotomas are present for a long period of time (several years) and are thus most likely associated with long-term adaptive processes, which cannot be simulated. Second, natural scotomas move with the eyes, whereas the AS in our experiment were locked to a particular location on the screen. Given the requirement to fixate, this may not have been a major issue. Third, we simulated the scotomas using smoothed edges, yet this is only an approximation as the actual transition zone between the blind and seeing field of patients is unknown. Incorporating prior knowledge on the presence of the scotomas has been shown previously to reduce pRF bias ^11,19^. Here, we show that the biases can be further reduced by using MP. However, the incorporation of an SS model with the purpose to detect visual field defects is redundant in the sense that it cannot be applied in clinical practice, given that the location of actual scotoma would not be unknown.

Due to the advanced age of the glaucoma participants, the presence of cognitive deficits was screened by means of a questionnaire. Only participants that reported no signs of depression, neurological, or psychiatric disorders were included in the study. However a deep screening of the participants cognitive abilities was not performed. Nonetheless, all the participants understood the instructions and had been in the scanner previously, which is evident from their good performance during the fixation task.

Eye movements change the position of the visual stimulus on the retina, which can affect the quality of the pRF mapping and induce biases in the pRF estimated properties, most commonly an enlargement of pRFs ^35–37^. Consequently, differences between the SAP and the reconstructed visual maps in glaucoma participants may also potentially result either from unstable fixation, or off-center fixation due to the presence of the scotoma. While eye movements were not recorded during scanning, participants did engage in an attentional task, which they performed on average with 78% accuracy. This task consisted of a fixation cross which covered the entire screen, this helps the participants with central scotomas to interpolate the most likely position of the center of the screen. These factors, together with the overall good quality of the retinotopic maps suggests eye-movement related factors did not play an important role.

Although in this study we did not test specifically the reliability of the fMRI-based VF reconstruction approaches and the robustness to noise, in previous work we showed that MP is reliable and robust to noise by performing a test-retest analysis and simulations with different levels of noise, (Carvalho et al., 2020). Given that the VF reconstruction is purely based on the MP output, we can conclude that MP VF are reliable and robust to noise. To verify this for VF reconstruction using pRF mapping, similar analysis will have to be applied.

## Conclusion

FMRI-based reconstruction of the visual field using pRF-based modeling enables the evaluation of vision loss and provides details on the properties of the visual cortex. Although the accuracy of the fMRI-based techniques was validated using simulations, in participants with glaucoma there were notable differences between the VF assessment done with fMRI and SAP. However, in contrast to SAP, fMRI-based VF reconstruction provides an objective measure of the quality of the VF. Moreover, it may provide information on the neurodegeneration underlying the loss of visual input, on cortical reorganization, and on the presence of predictive masking. Finally we propose the use of fMRI-based VF assessments in a longitudinal study, to evaluate the technique’s feasibility to monitor disease progression and treatment response. This will be particularly important in patients unable to perform SAP.

## Acknowledgments

Authors JC and AI were supported by the European Union’s Horizon 2020 research and innovation programme under the Marie Sklodowska-Curie grant agreements No. 641805 (NextGenVis) and No. 661883 (EGRET). JM was supported by the European Union’s Erasmus + program. The funding organization had no role in the design, conduct, analysis, or publication of this research. ^38^

## Supplementary Information

### 1. Reliability of the performance of participants with Glaucoma during SAP

**Table S1.** Reliability of the performance of participants with glaucoma during SAP. Fixation losses, % of false positive and false negative error.

| Participants | HFA fixation losses | False Pos error (%) | False Neg error (%) |
| --- | --- | --- | --- |
| G01 (L/R) | 0/11 \| 1/10 | 0 \| 0 | 0 \| 0 |
| G02 (L/R) | 2/12 \| 1/13 | 1\|4 | 3\|5 |
| G03 (L/R) | 0/13 \|1/14 | 0\|2 | 0\|7 |
| G04 (L/R) | 0/13 \| 1/14 | 0\|0 | 0\|5 |
| G05 (L/R) | 0/12 \| 1/10 | 2\|1 | 3\|4 |
| G06 (L/R) | 4/11 \| 1/11 | 0\|4 | 1\|0 |
| G07 (L/R) | 3/15 \| 6/13 | 3\|7 | 9\|3 |
| G08 (L/R) | 0/11 \| 1/10 | 0\|3 | 6\|3 |
| G09 (L/R) | 0/16 \| 0/17 | 1\|2 | 0\|7 |
| G10 (L/R) | 3/10 \| 1/11 | 0\|0 | 0\|0 |
| G11 (L/R) | 0/12 \| 0/15 | 4\|1 | 2\|7 |
| G12 (L/R) | 2/14 \| 2/13 | 7\|0 | 0\|0 |
| G13 (L/R) | 0/15 \| 1/12 | 0\|7 | 0\|9 |
| G14 (L/R) | 0/11 \| 0/14 | 4\|0 | 0\|6 |
| G15 (L/R) | 0/12 \| 1/12 | 5\|0 | 6\|1 |
| G16 (L/R) | 0/13 \| 1/14 | 4\|1 | 12\|6 |
| G17 (L/R) | 0/15 \| 0/12 | 0\|0 | 5\|9 |
| G18 (L/R) | 0/11 \| 1/12 | 0\|0 | 3\|0 |
| G19 (L/R) | 1/13 \| 0/12 | 0\|3 | 0\|7 |

### 2. Selection of number of probes

The number of probes to be included in the probe map was selected empirically. Using simulations probe maps were calculated with several numbers of probes and the time to compute was recorded (Table S1). The optimal number of probes represents a trade off between: 1) ensuring that the entire field of view is sampled at a fine scale and 2) the time that it takes to compute. Figure S1 shows the probe maps using different number of probes calculated based on simulated time series of a pRF located at x=3,y-3 and size=1 deg. To make the simulations more realistic artificial noise was added (SNR=2). The method used to generate the simulated time series is described in ^26^. Figure and table S1 shows that using 10000 probes results in a good coverage of the entire visual field within a reasonable time.

**Figure S1.**
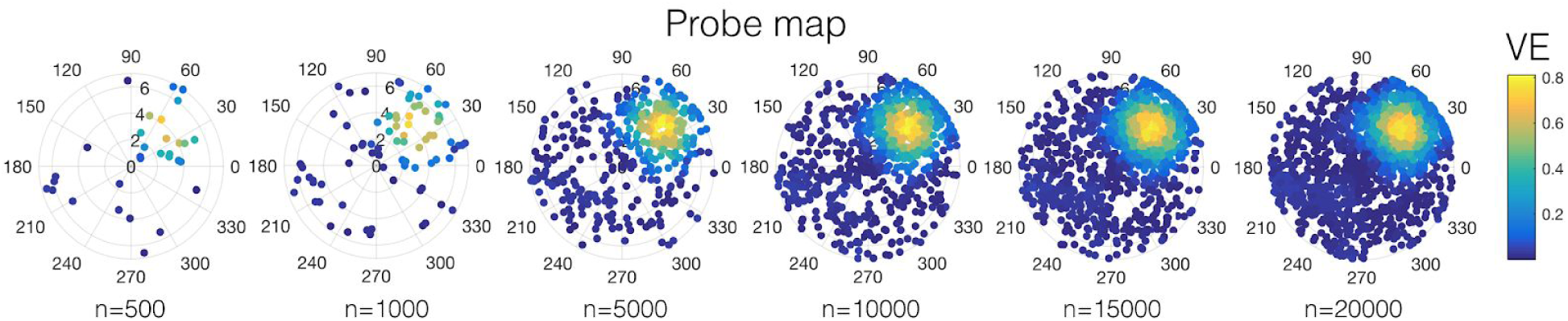
Probe maps calculated with different number of probes. The colormap represents the variance explained.

**Table S2.** Average computation time per voxel using different numbers of probes. The time was calculated using matlab 2018b on a macbook pro.

| Number of probes | Average computation time per voxel (s) |
| --- | --- |
| 500 | 1.2 |
| 1000 | 2.5 |
| 5000 | 12.8 |
| 10000 | 22.6 |
| 15000 | 34.2 |
| 20000 | 46.7 |

**Table S3.**
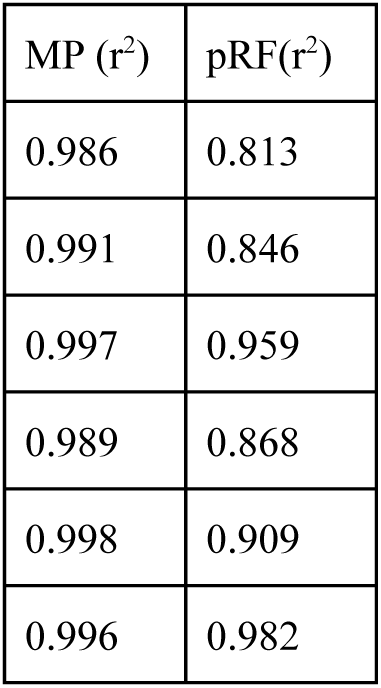
Correlation coefficient of the spared VF (region of the VF without SS) between SF and FF models for MP and pRF approaches.

**Figure S2.**
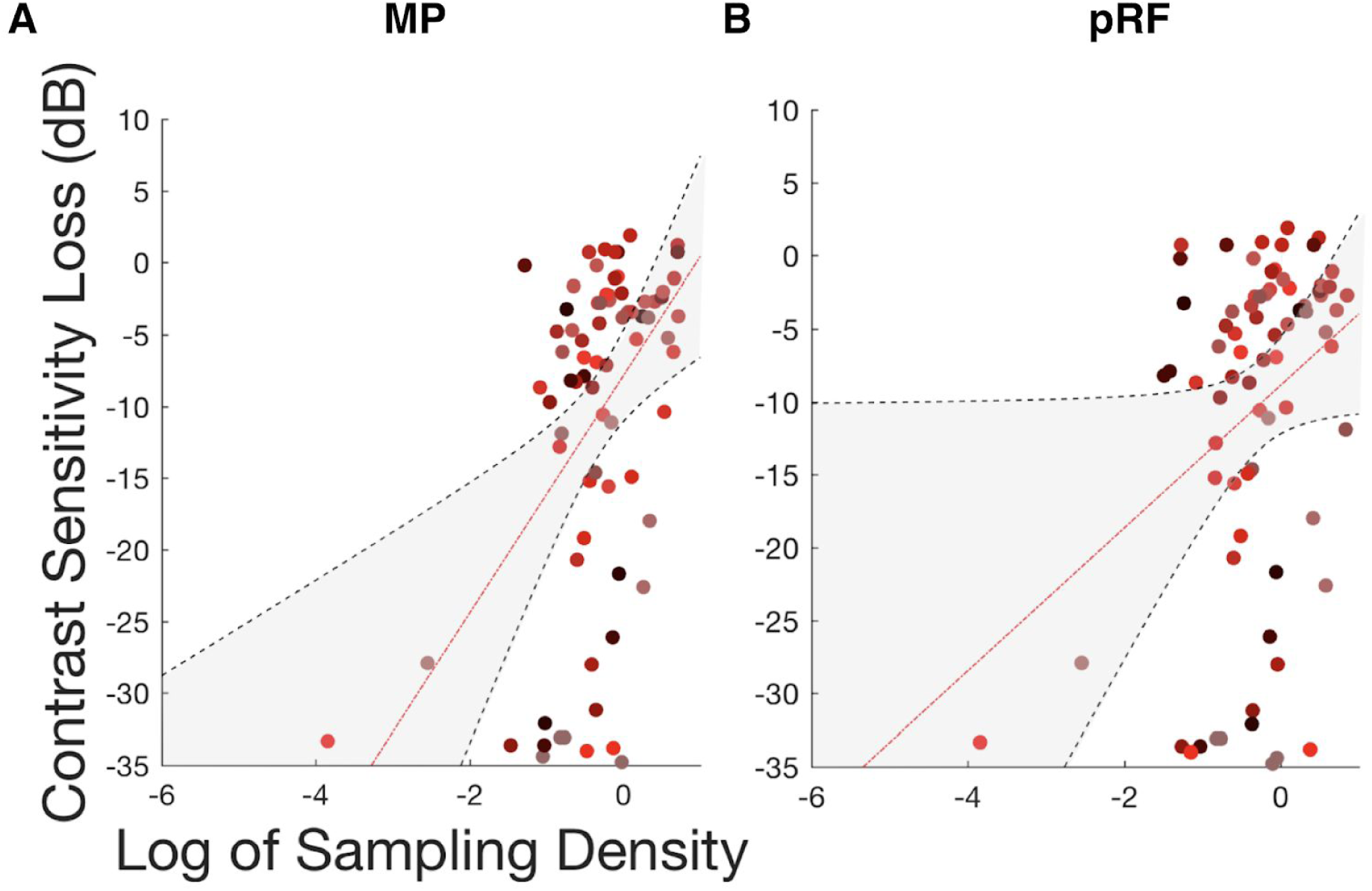
Correlation of sampling density measured with fMRI-based techniques and contrast sensitivity loss obtained with SAP. Panels A and B show the correlation of the sampling density of participants with glaucoma obtained from individual quadrants with contrast sensitivity loss, for MP (A) and pRF (B) techniques, respectively. Each data point is from a separate quadrant of an individual participant with glaucoma. Each color represents the data point of a single participant. The dashed red line represents the linear fit to the data and the shaded region represents the 95% confidence interval of the fitted parameters. Note that the correlations were obtained using the same visual field area for the fMRI-based visual sampling and the SAP-based contrast sensitivity (a 7×7 deg quadrant adjacent to fixation).

